## supplementary material for "Separability and Geometry of Object Manifolds in Deep Neural Networks"

February 1, 2020

### 1 Figures

#### 1.1 Object manifolds in deep convolutional networks (AlexNet, VGG-16, ResNet-50)

In this section, we demonstrate that the overall trend of improvement in capacity, reduction in dimension and radius across the layers of a deep network generalizes to additional data-sets. Those include additional manifolds and a family of network models, residual networks, from which we focus on ResNet-50. We show the results on AlexNet, VGG-16 alongside ResNet-50 for comparison.

##### 1.1.1 Capacity for point-cloud manifolds in AlexNet, VGG-16, ResNet-50

The capacity of point-cloud manifolds created from ImageNet classes increases across the layers of ResNet-50 (figure 1). Furthermore, the capacity measured in AlexNet, VGG-16, ResNet-50 exhibit the same trend as their respective performance in the ImageNet object classification task for which they were trained. This trend continues when using residual networks of different depths (figure 2a), where most of the capacity improvement is achieved in the final layers.

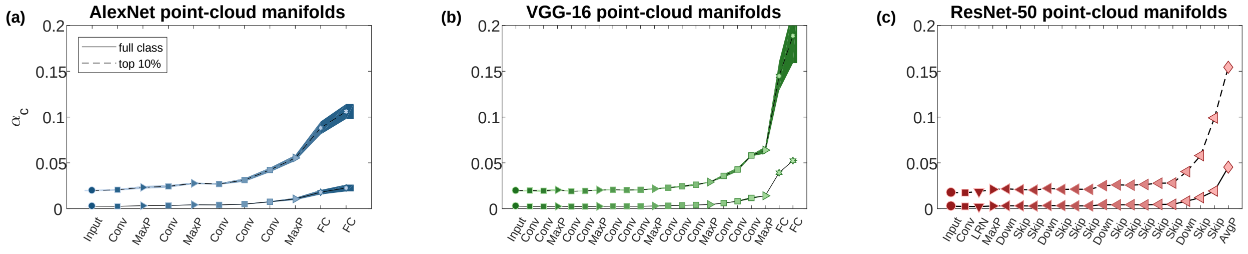

**Figure 1: Capacity for point-cloud manifolds created from ImageNet classes.** Classification capacity for point-cloud manifolds (full line: full class manifolds; dashed line: top 10% manifolds, see main text Methods) along the layers of deep neural networks: AlexNet (a) VGG-16 (b) and ResNet-50 (c). Line and markers at (a,b) indicate mean value over five choices of 50 objects and surrounding shaded areas indicate 95% confidence interval, while for (c) only the first set of objects is used.
The x-axis labels provides abbreviation of the layer types ('Input'- pixel layer, 'Conv'- convolutional layer, 'MaxP'- max-pooling layer, 'FC'- fully connected layer, 'AveP'- average pooling layer, 'LRN'- local normalization layer, 'Skip'- skip module, 'Down'- skip module with downsampling). Marker shape represents layer type (circle- pixel layer, square- convolution layer, right-triangle- max-pooling layer, hexagon- fully connected layer, diamond- average pooling layer, down-triangle- local normalization layer, left-triangle- a skip module). Color (blue- AlexNet, green- VGG-16, red- ResNet-50) changes from dark to light along the network layers. Features in linear layers ('Conv', 'FC') and skip modules ('Skip', 'Down') are extracted after a ReLU non-linearity.

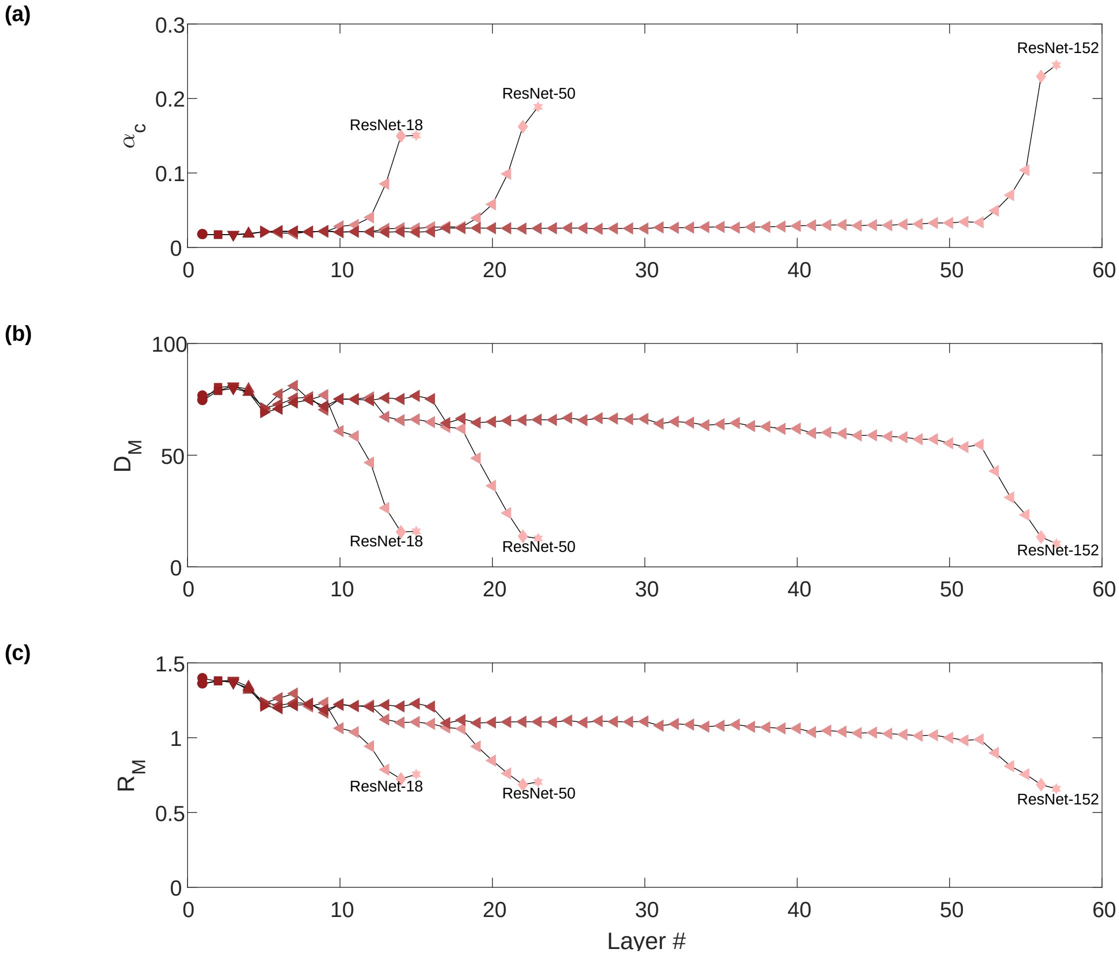

**Figure 2: Capacity and geometry for point-cloud manifolds using residual networks.**
Classification capacity (a) mean manifold dimensions (b) and mean manifold radii (c) for point-cloud top 10% manifolds along the layers of deep residual networks (ResNet-18, ResNet-50, ResNet-152).
ResNet-50 results are also reported in other supplementary figures. The x-axis indicates layer depth starting from the pixel layer. Marker shape represents layer type (circle- pixel layer, square- convolution layer, right-triangle- max-pooling layer, hexagon- fully connected layer, diamond- average pooling layer, down-triangle- local normalization layer, left-triangle- a skip module). Features in linear layers and skip modules are extracted after a ReLU non-linearity.

##### 1.1.2 Capacity for smooth manifolds in AlexNet, VGG-16, ResNet-50

When calculating capacity for smooth manifolds, a finite number of samples is used; figure 3 justify this practice by showing that for both 1-d and 2-d smooth manifolds when the number of samples tends to infinity capacity is still finite, and is very closed to the values measured using the maximal number of samples.

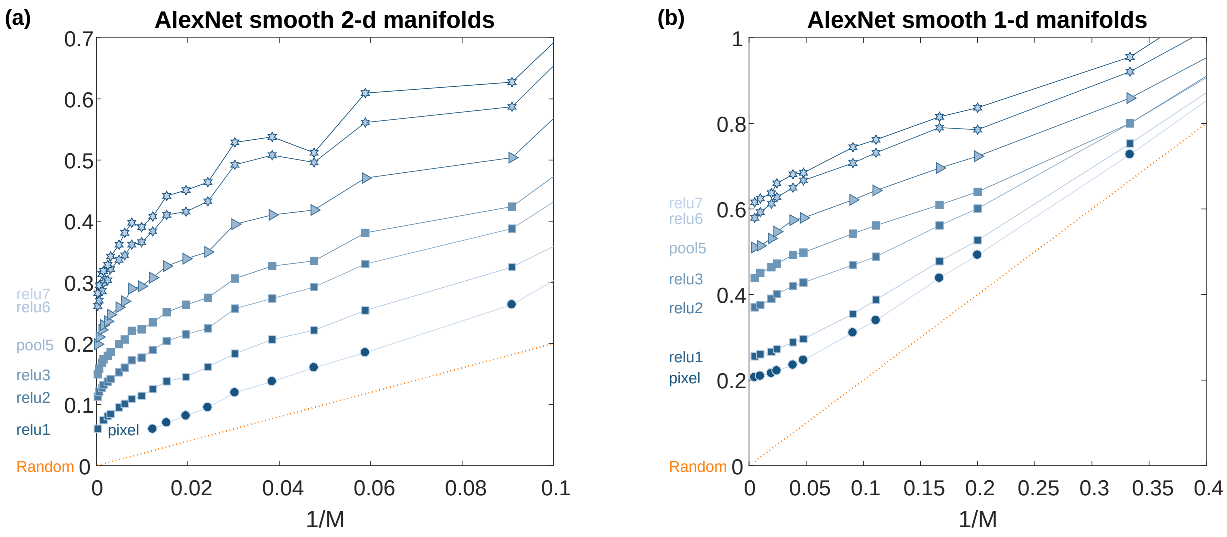

**Figure 3: Capacity dependence on the number of samples in smooth manifolds.**
Numerically measured capacity for smooth manifolds across the layers of AlexNet using different number of samples (M) per manifold. The orange dashed line indicates the capacity expected for random points (2/M, see main text Methods).
(a) capacity (y-axis) vs. inverse the number of samples (x-axis) for 2-d smooth manifolds.
(b) capacity (y-axis) vs. inverse the number of samples (x-axis) for 1-d smooth manifolds.
Marker shape represents layer type (circle- pixel layer, square- convolution layer, right-triangle- max-pooling layer, hexagon- fully connected layer). Color changes from dark to light along the network.

The capacity of smooth manifolds created from warped ImageNet images increases across the layers of ResNet-50 (figure 4), as demonstrated for AlexNet and VGG-16 at main figure 5. As in that figure, the capacity improvement increases supra-linearly with the complexity of the manifold, quantified by its total variability at the pixel layer.

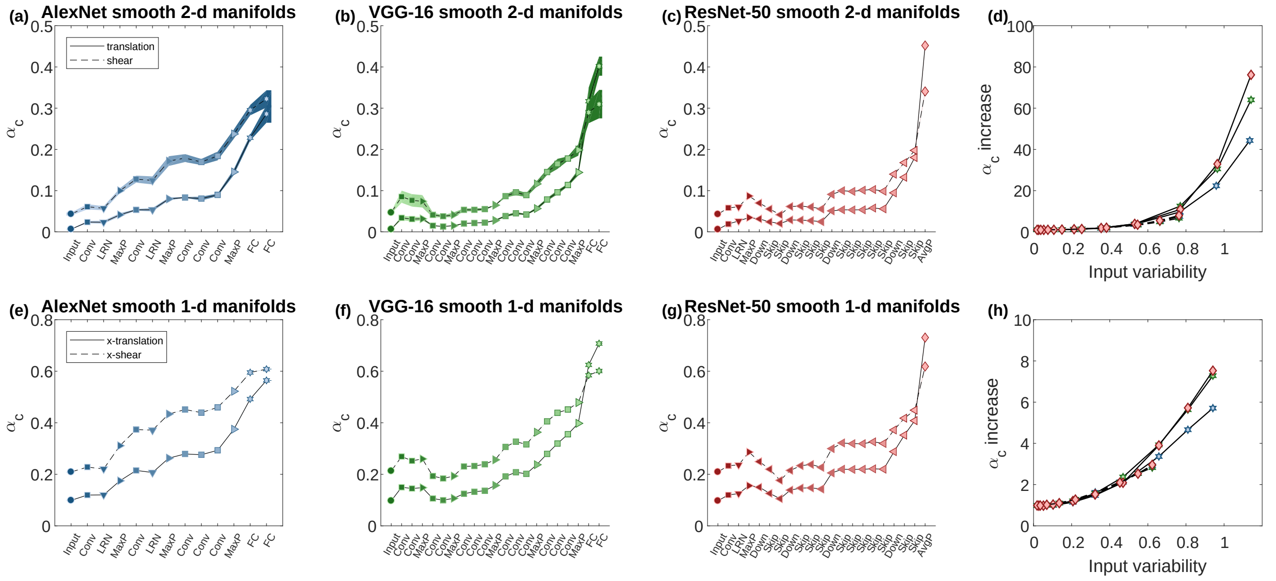

**Figure 4: Capacity for smooth manifolds created from warped ImageNet images.**
(a-c,e-g) Classification capacity for smooth manifolds (full line: translation; dashed line: shear) along the layers of AlexNet (a,e), VGG-16 (b,f) and ResNet-50 (c,g). The x-axis labels provide an abbreviation of the layer types. Line and markers at (a,b) indicate mean value over four choices of 64 objects and surrounding shaded areas indicate 95% confidence interval, while on other panels, only the first set of objects is used. Marker shape represents layer type (circle- pixel layer, square- convolution layer, right-triangle- max-pooling layer, hexagon- fully connected layer, diamond- average pooling layer, down-triangle- local normalization layer, left-triangle- a skip module). Features in linear layers and skip modules are extracted after a ReLU non-linearity. Color (blue- AlexNet, green- VGG-16, red- ResNet-50) changes from dark to light along the network.
(d,h) Capacity increase from the input (pixel layer) to the output (features layer) of AlexNet (blue markers), VGG-16 (green markers) and ResNet-50 (red markers) for smooth manifolds created from 2-d translation (d) or 1-d translation (h). The capacity increase is the ratio of capacity at the features layer and the pixel layer (y-axis), at different levels of stimuli variation, measured using supplementary equation (3) at the pixel layer (x-axis).

##### 1.1.3 Geometry of point-cloud and smooth manifolds in AlexNet, VGG-16, ResNet-50

The geometry of point-cloud and smooth manifolds is characterized by the manifolds' dimensions and radii.

The mean values of manifold dimensions, across the layers of AlexNet, VGG-16, and ResNet-50, are shown in figure 5a-f, along with 95% confidence intervals with respect to the sampling of different objects. This serves to demonstrate a large but non-monotonic reduction in dimension along each hierarchy. Going beyond the mean, the distribution of dimension values is shown for selected manifolds in figure 5g-i. The manifold dimension $D_M$ is comparable in magnitude and layer-by-layer changes to a well-known measure of dimensionality, the spectral participation ratio $D_{svd} = (\Sigma_i \lambda_i^2)^2/(\Sigma_i \lambda_i^4)$ where $\{\lambda_i\}_{i=1}^M$ are the manifold spectrum (i.e. singular values of centered manifold samples), shown for comparison at figure 5j-l.

Similarly, the mean values of manifold radii, along the layers of AlexNet, VGG-16 and ResNet-50, are shown in figure 6a-f. Those demonstrate a small and monotonic reduction in radius along each hierarchy for point-cloud manifolds, while for smooth manifolds, the main decrease is in the first layer and last layers (see discussion in the main text). Going beyond the mean, the distribution of radii values is shown for selected manifolds in figure 6g-i.

For residual networks of different depths, we find that the improvement of capacity in the final layers is associated with a decrease of both radius and dimension (figure 2b-c).

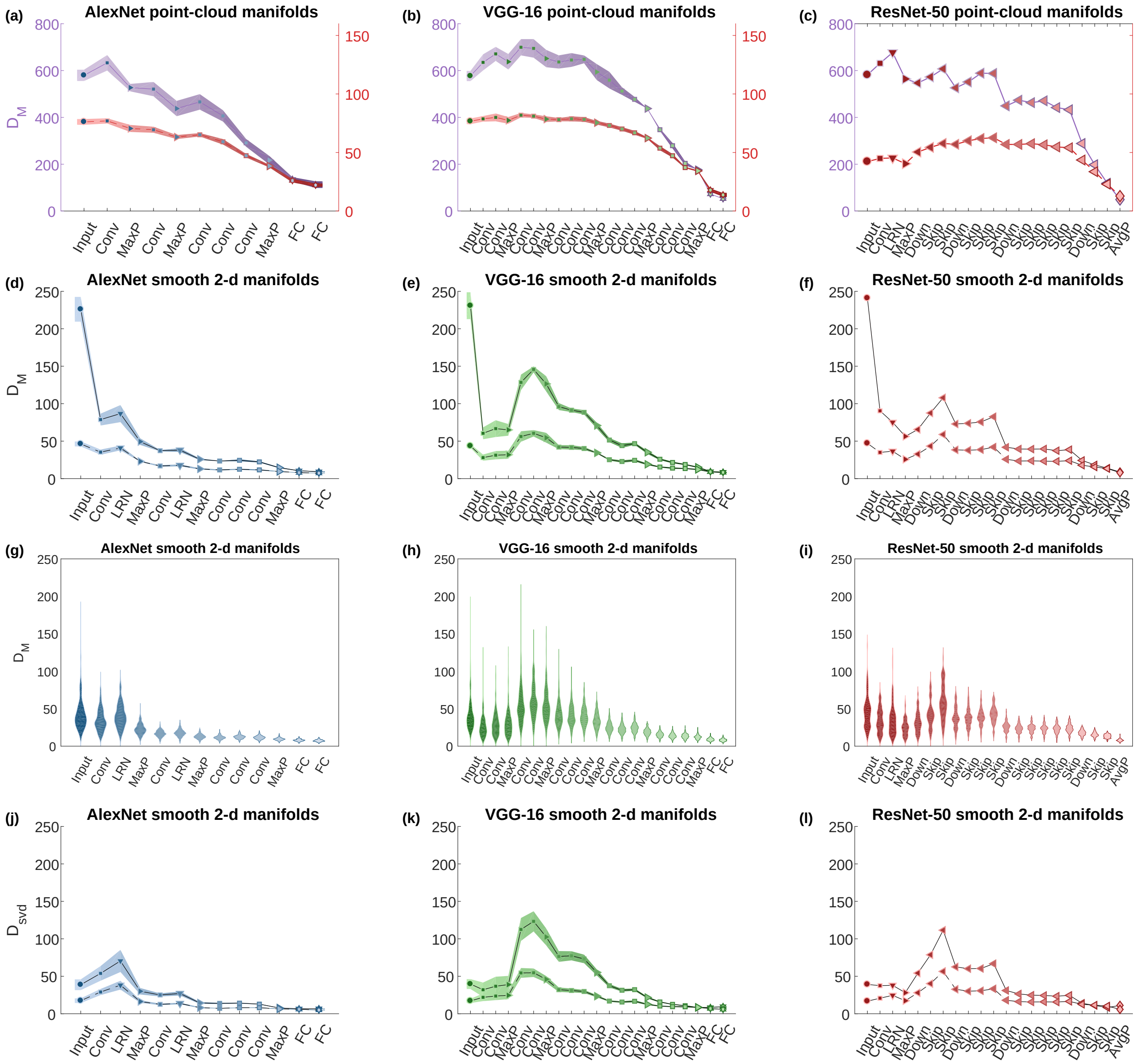

**Figure 5: Manifold dimensions along the hierarchy.**
(a-c) Mean manifold dimension for point-cloud manifolds of AlexNet (a), VGG-16 (b) and ResNet-50 (c); full-line: full-class manifolds, dashed-line: top 10% manifolds. Values of top 10% manifolds are shown against a secondary y-axis (color-coded by the edge of the marker) to improve visibility. (d-f) Mean manifold dimension for smooth 2-d manifolds of AlexNet (d), VGG-16 (e) and ResNet-50 (f); full-line: translation manifolds, dashed-line: shear manifolds. (g-i) Distribution of manifold dimension for smooth 2-d shear manifolds for the same deep networks as (d-f), illustrated as a per-layer histogram (kernel width is std/5). (j-l) Mean manifold participation ratio for smooth 2-d manifolds of AlexNet (j), VGG-16 (k) and ResNet-50 (l); full-line: translation manifolds, dashed-line: shear manifolds.
Line and markers at (a,b,d,e,j,k) indicate mean value over different choices of objects and surrounding shaded areas indicate 95% confidence interval, while at (c,f,l) only the first set of objects is used. The x-axis labels provide an abbreviation of the layer types. Marker shape represents layer type (circle- pixel layer, square- convolution layer, right-triangle- max-pooling layer, hexagon- fully connected layer, diamond- average pooling layer, down-triangle- local normalization layer, left-triangle- a skip module). Features in linear layers are extracted after a ReLU non-linearity. Color (blue- AlexNet, green- VGG-16, red- ResNet-50) changes from dark to light along the network.

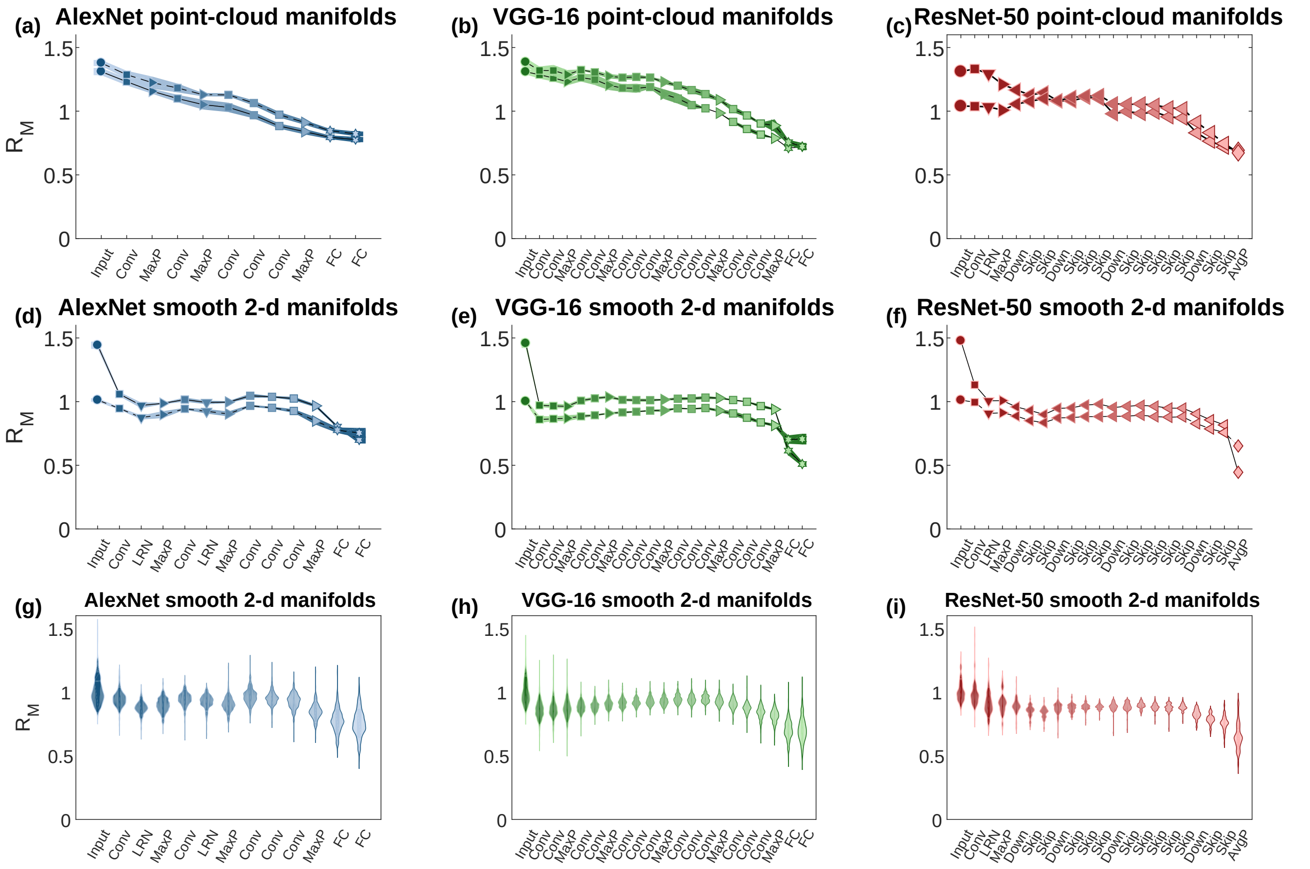

**Figure 6: Manifold radii along the hierarchy.**
(a-c) Mean manifold radius for point-cloud manifolds of AlexNet (a), VGG-16 (b) and ResNet-50 (c); full-line: full-class manifolds, dashed-line: top 10% manifolds. (d-f) Mean manifold radius for smooth 2-d manifolds of AlexNet (d), VGG-16 (e) and ResNet-50 (f); full-line: translation manifolds, dashed-line: shear manifolds. (g-i) Distribution of manifold radius for smooth 2-d shear manifolds for the same deep networks as (d-f), illustrated as per-layer histogram (kernel width is std/5).
Line and markers at (a,b,d,e) indicate mean value over different choices of objects and surrounding shaded areas indicate 95% confidence interval, while at (c,f) only the first set of objects is used. The x-axis labels provide an abbreviation of the layer types. Marker shape represents layer type (circle- pixel layer, square- convolution layer, right-triangle- max-pooling layer, hexagon- fully connected layer, diamond- average pooling layer, down-triangle- local normalization layer, left-triangle- a skip module). Features in linear layers are extracted after a ReLU non-linearity. Color (blue- AlexNet, green- VGG-16, red- ResNet-50) changes from dark to light along the network.

Using the theoretical relation between capacity and manifold properties (demonstrated empirically at main figure 10) figure 7 demonstrates that most of the observed changes in capacity can be attributed to changes in dimensionality while changes in radii contribute just in the final layers. When manifolds radii are fixed at their pixel layer values, a large fraction of the capacity improvement would still occur, but when manifolds dimensions are fixed at their pixel layer values capacity would hardly improve.

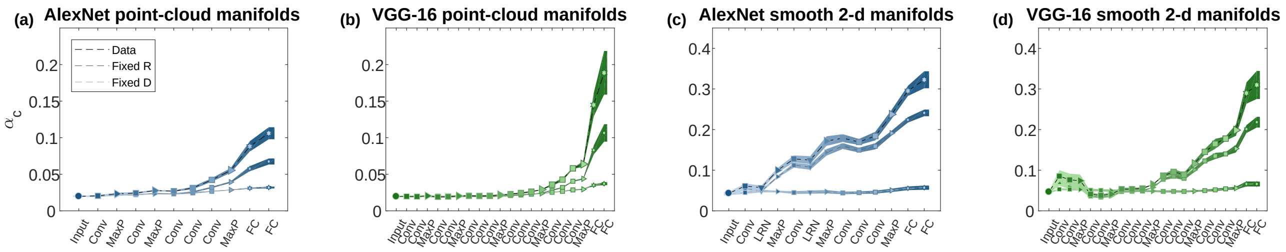

**Figure 7: Relation between changes of capacity and changes of radii and dimensions.** Classification capacity for point-cloud manifolds (top 10%) along the layers of AlexNet (a), VGG-16 (b) and for smooth 2-d shear manifolds along the layers of AlexNet (c), VGG-16 (d). The results using the real data (black lines), previously presented in main figures 4 and 5, are to be compared to the expected capacity with the observed dimensions but the radii fixed at their values in the pixel layer (dark gray lines), or the expected capacity with the observed radii but the dimensions fixed at their values in the pixel layer (light gray lines). Shaded areas represent 95% confidence intervals with respect to the sampling of different objects.
The x-axis labels provide an abbreviation of the layer types. Marker shape represents layer type (circle- pixel layer, square- convolution layer, right-triangle- max-pooling layer, hexagon- fully connected layer, down-triangle- local normalization layer). Features in linear layers are extracted after a ReLU non-linearity. Color (blue- AlexNet, green- VGG-16) changes from dark to light along the network.

##### 1.1.4 Geometry of 1-d versus 2-d smooth manifolds in AlexNet, VGG-16, ResNet-50

To quantify the complexity of internal representations at smooth manifolds, we measure a fraction between manifold dimensions created by 2-d variation and 1-d variation in the latent parameter space (figure 8a-c). When $D_M^{2d}/D_M^{1d}$ is 2, it means that the dimensionality (in neural space) of 2-d variation is exactly twice the dimensionality created by 1-d variation. This can be either because the subspaces created by 2-d variation are factorized, or because the created manifold is small. If $D_M^{2d}/D_M^{1d}$ is large (above 2), that means that there is a nonlinear interaction between the dimensions created by the 2-d variations in the latent space, and additional dimensions are created as a result. Here we find this is the case when manifold radii are large enough, as evident for translation manifolds which are very high-dimensional at the pixel layer. On the other hand, $R_M^{2d}/R_M^{1d}$ is 1 throughout this data-set, as expected when the variability induced by each of 1-d transformation is arranged in orthogonal axes (figure 8d-f).

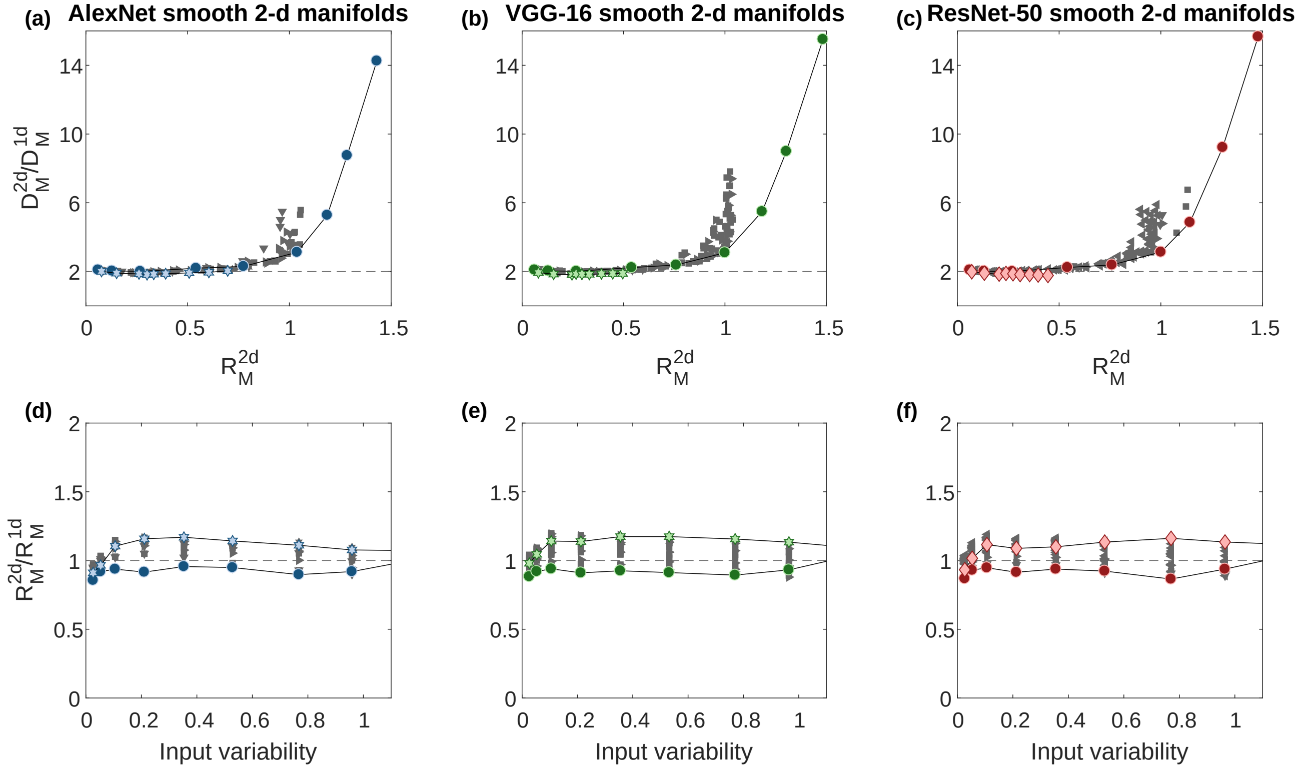

**Figure 8: Comparison of the geometry of 2-d and 1-d manifolds.**
(a-c) Manifolds dimensions for smooth 2-d manifolds using AlexNet (a), VGG-16 (b) and ResNet-50 (c) relative to the average dimensions of the corresponding 1-d manifolds (i.e., the two 1-d manifolds with the same maximal object displacement) at the y-axis, shown against the 2-d manifolds radii at the x-axis.
(d-f) Manifolds radii for smooth 2-d manifolds using AlexNet (d), VGG-16 (e) and ResNet-50 (f) relative to the average radii of the corresponding 1-d manifolds (y-axis), shown against the 2-d stimuli variability, measured using supplementary equation (3) at the pixel layer (x-axis).
Marker shape represents layer type (circle- pixel layer, square- convolution layer, right-triangle- max-pooling layer, hexagon- fully connected layer, diamond- average pooling layer, down-triangle- local normalization layer, left-triangle- a skip module). Only results from the first and last layers are colored; other layers are in gray to avoid clutter.

##### 1.1.5 Manifold correlations and their effect on capacity in AlexNet, VGG-16, ResNet-50

Here we demonstrate that different types of between-manifold correlations (i.e. center-center, axis-axis) are reduced across the layers of the deep hierarchies analyzed, for both point-cloud and smooth manifolds (figure 9). Consider $P$ manifolds which correspond to the responses of $N$ neurons to $M$ samples, denoted $F^{\mu}_{i,m}$ for $\mu = 1..P$, $i = 1..N$ and $m = 1..M$. Each manifold is described by its center $\vec{x}^{\mu}$ and its singular value decomposition (SVD) $F^{\mu}_{i,m} = x^{\mu}_i + \Sigma_l \lambda^{\mu}_l u^{\mu,l}_i v^{\mu,l}_m$ for non-negative scalars $\{\lambda^{\mu}_l\}_l$ and orthonormal sets of vectors $\{\vec{u}^{\mu,l} \in \mathbb{R}^N\}_l, \{\vec{v}^{\mu,l} \in \mathbb{R}^M\}_l$.
Then center correlations are defined

$$\rho_{CC} = \left\langle \frac{|\vec{x}^{\mu} \cdot \vec{x}^{\nu}|}{||\vec{x}^{\mu}|| \cdot ||\vec{x}^{\nu}||} \right\rangle_{\mu \neq \nu} \tag{1}$$

and axes correlations are defined

$$\rho_{AA} = \left\langle \sum_l \hat{\lambda}^{\mu}_l \hat{\lambda}^{\nu}_l |\vec{u}^{\mu,l} \cdot \vec{u}^{\nu,l}| \right\rangle_{\mu \neq \nu} \tag{2}$$

where $\hat{\lambda}^{\mu}_l = \lambda^{\mu}_l / \sqrt{\sum_l (\lambda^{\mu}_l)^2}$. Furthermore, the total manifold variability is defined:

$$v = \left\langle \frac{\sqrt{\sum_l (\lambda^{\mu}_l)^2}}{||\vec{x}^{\mu}||} \right\rangle_{\mu} \tag{3}$$

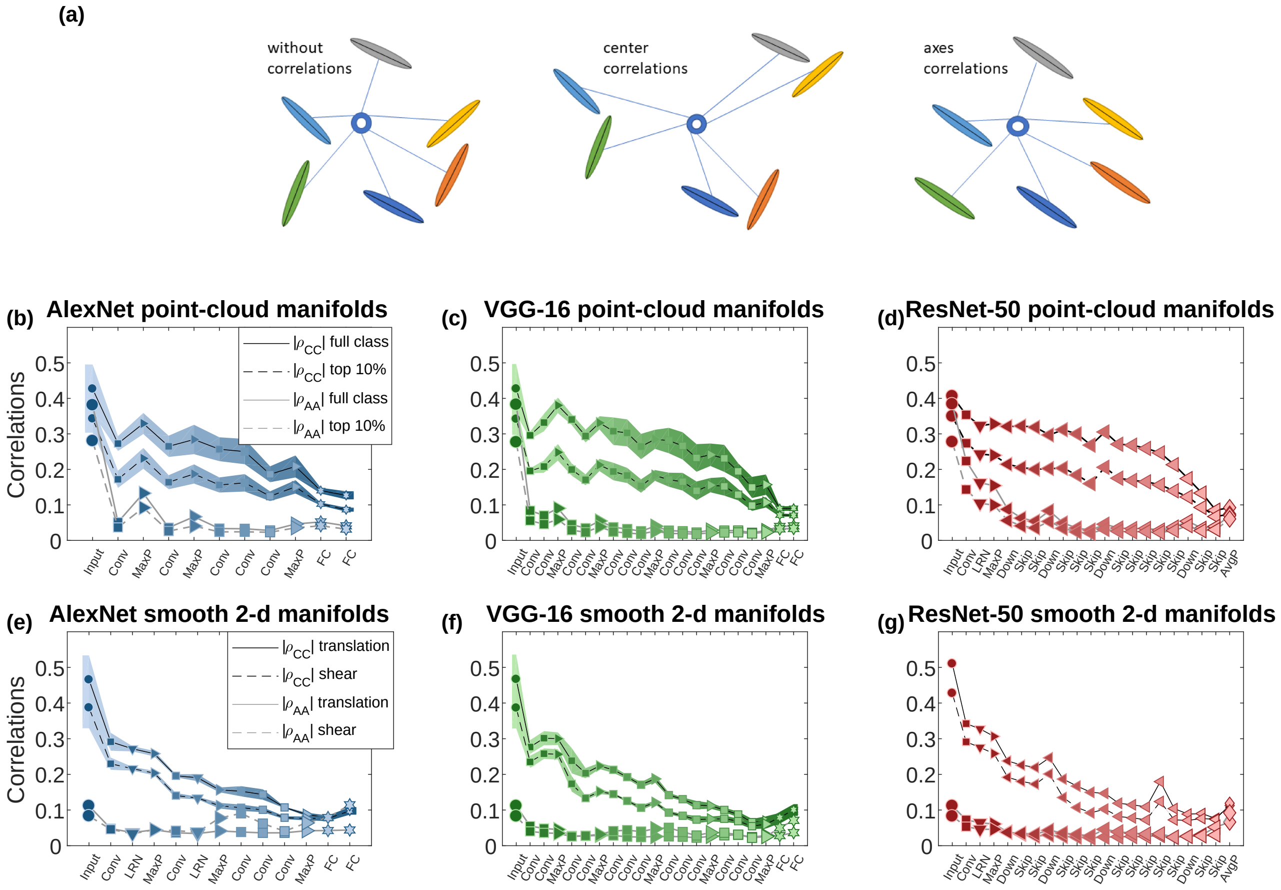

**Figure 9: Center and axes correlations between manifolds.** (a) illustration of object manifolds without correlations (left), where the manifolds' centers are random points and the manifolds' main axes of variation are randomly oriented; with center correlations (middle), where the manifolds' centers are clustered; with axes correlations (right), where the manifolds' main axes of variation are aligned.
(b-d) Center and axes correlations for point-cloud manifolds using AlexNet (b), VGG-16 (c) and ResNet-50 (d); black- center correlations; gray- axes correlations; full line- full class manifolds, dashed line- top 10% manifolds.
(e-g) Center and axes correlations for smooth manifolds using AlexNet (e), VGG-16 (f) and ResNet-50 (g); black- center correlations; gray- axes correlations; full line- translation manifolds, dashed line- shear manifolds.
For center correlations of AlexNet and VGG-16 line and markers indicate mean value over different choices of objects; surrounding shaded areas indicate 95% confidence interval, with some of the results previously presented as "after training" results of main figure 7. For axes correlations, ResNet-50 only the first set of objects was used.
The x-axis labels provide an abbreviation of the layer types. Marker shape represents layer type (circle- pixel layer, square- convolution layer, right-triangle- max-pooling layer, hexagon- fully connected layer, diamond- average pooling layer, down-triangle- local normalization layer, left-triangle- a skip module). Features in linear layers and skip modules are extracted after a ReLU non-linearity. Color (blue- AlexNet, green- VGG-16, red- ResNet-50) changes from dark to light along the network.

Furthermore, comparing capacity from the data manifolds with capacity on surrogate data manifolds where different types of correlations are removed through randomization allows us to quantify those center correlations have a substantial effect while axes correlations have a much smaller effect (figure 10). As predicted by theory, center-correlations decrease capacity while axes correlations increase capacity, relative to a correlations-free baseline.

The creation of data without center correlations is done by replacing the center of each manifold with a random vector. The creation of data without axes correlations is done by applying a random permutation to the neurons of each manifold. Then by comparing numerically calculated capacity (see Methods in the main text) from the original data to that calculated on the surrogate data, the effect of the removed correlations is revealed.

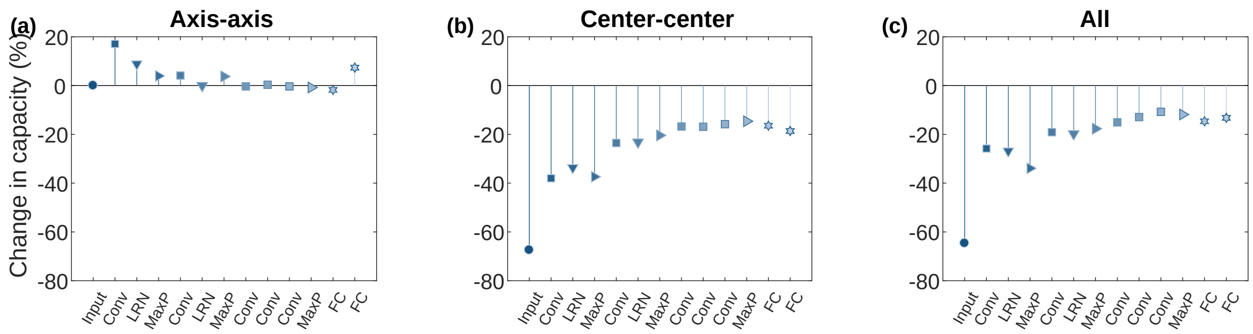

**Figure 10: Quantifying correlations' effects on capacity through randomization.**
Change of numerically measured capacity due to a randomization procedure that produces manifolds where different kinds of correlations are absent. Results are shown as percent changes of original data capacity relative to the capacity of the randomized data. All results are calculated from smooth 1-d shear manifolds.
(a) The effect of removing axes correlations at different layers of AlexNet.
(b) The effect of removing center correlations at different layers of AlexNet.
(c) The effect of removing both correlations at different layers of AlexNet.
The x-axis labels provide an abbreviation of the layer types. Marker shape represents layer type (circle- pixel layer, square- convolution layer, right-triangle- max-pooling layer, hexagon- fully connected layer, down-triangle- local normalization layer). Features in linear layers are extracted after a ReLU non-linearity. Color changes from dark to light along the network.

###### 1.1.6 Deep network building-blocks have a consistent effect on manifold geometry and correlations

As discussed in the main text, the role of different network building-blocks is explored by analyzing the effect on the capacity of single operations and computational building-blocks, i.e., common operation sequences used in DCNNs (figure 8 in the main text). Figure 11 complements this discussion by exhibiting results for linear operations (i.e., convolutional and fully-connected layers) and partial building blocks (convolution followed by ReLU, without terminating max-pooling operation as in the main text). In both cases, the changes due to those building blocks are much less consistent than that of the full sequence.

Additionally, a similar analysis of the effect of a "skip module", a sequence of operations used in "Residual Networks", reveal that those modules usually reduce the dimension, radius, and correlations, with a clear trade-off effect (cases where dimension is not reduced, are associated with reduction of correlations and vice-versa). Indeed skip modules replace in those networks the use of convolutions followed by ReLU and max-pooling operations [1].

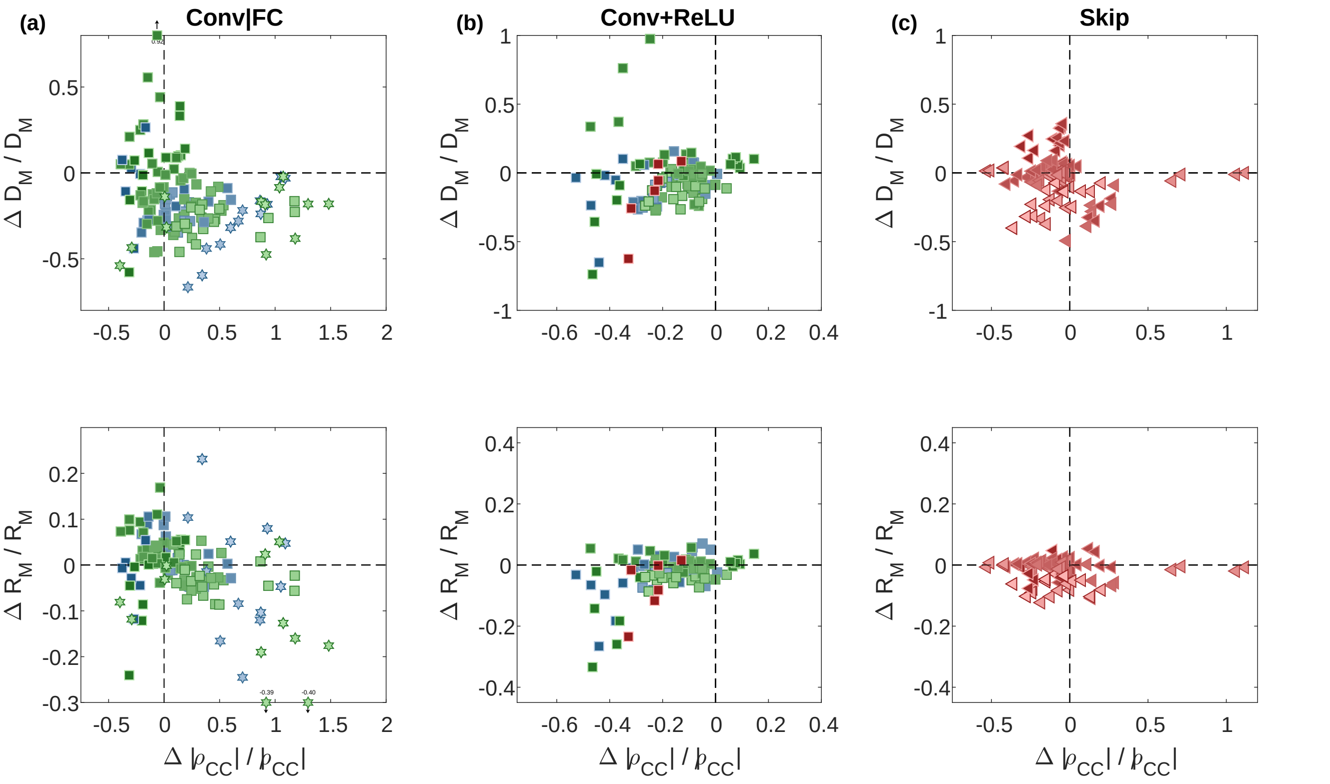

**Figure 11: Manifold property changes by network building blocks.** Changes in the relative manifold properties between the input and the output of different network building-blocks, shown as a change in dimension vs. change in center correlations (top) and change in radius vs. change in center correlations (bottom). Each panel pool results from a specific building-block in AlexNet (blue markers), VGG-16 (green markers) and ResNet-50 (red markers) for both point-cloud manifolds (full class, top 10%) and smooth manifolds (1-d and 2-d, translation and shear).
(a) Changes in manifold properties for isolated linear operations (i.e., convolution, FC).
(b) Changes in manifold properties for a common sequence of layers, convolution followed by ReLU operation.
(c) Changes in manifold properties for "skip modules", a common sequence in the ResNet-50 architecture.
Marker shape represents layer type (square- convolution layer, hexagon- fully connected layer, left-triangle- a skip module). For sequences, a marker that matches the type of the first operation is used. Color changes from dark to light along the network.

#### 1.2 Predictions of manifold separability theory

In this section we demonstrate that the theory of linear separability of manifolds provides interesting predictions that may be verified for manifolds created by the layers of DCNNs.

##### 1.2.1 Comparison between theory and numerically measured capacity in smooth manifolds of AlexNet, VGG-16, ResNet-50

Here we demonstrate that the manifolds classification capacity computed from the full mean-field theory (main text equation (1)) matches capacity measured numerically (see Methods in the main text). Figure 12 presents an excellent match across all layers and manifold variability levels using different DCNNs.

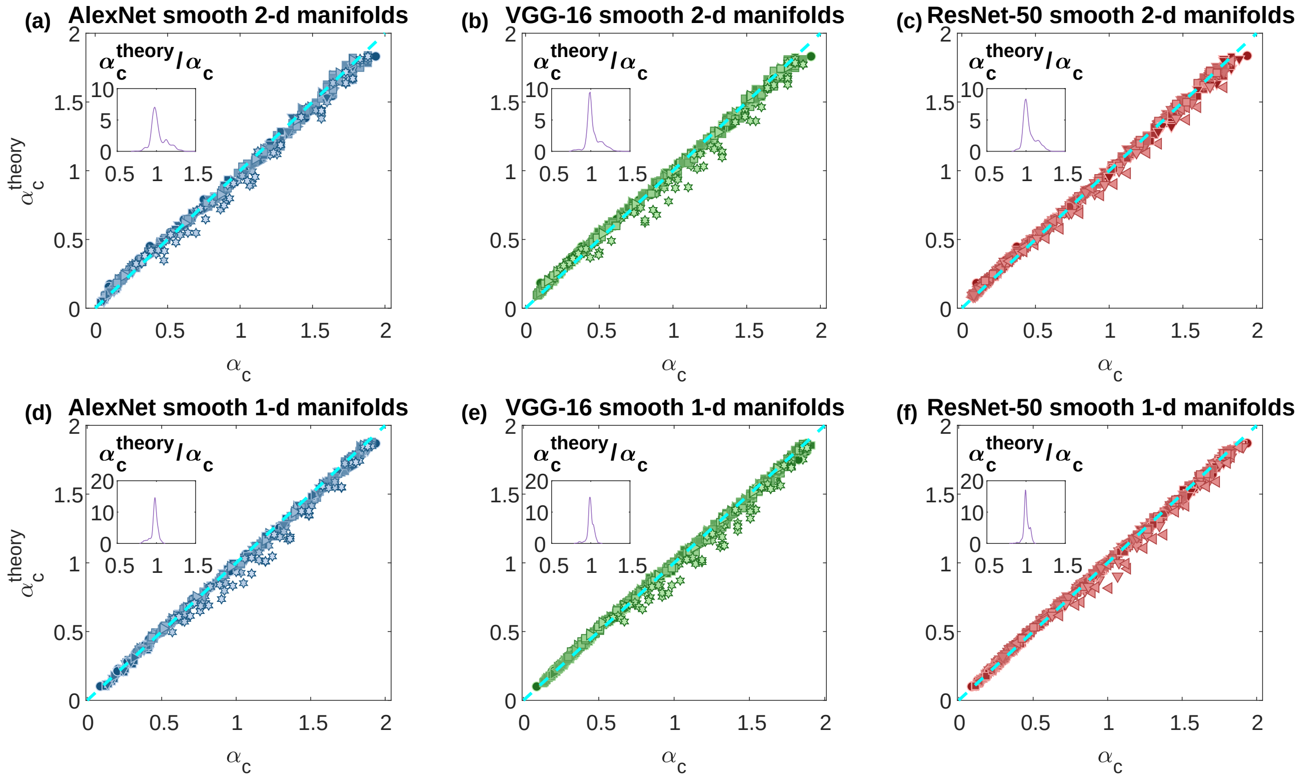

**Figure 12: Comparison of capacity measured using theory and numerically.**
Comparison of numerically measured capacity (x-axis) with the theoretical prediction (y-axis) for smooth manifolds at different layers along the hierarchy and different manifold variability levels. Inset: histogram of the ratio between the y-axis and the x-axis.
(a-c) Results for smooth 2-d manifolds across AlexNet (a), VGG-16 (b) and ResNet-50 (c)
(d-f) Results for smooth 1-d manifolds across AlexNet (d), VGG-16 (e) and ResNet-50 (f)
Marker shape represents layer type (circle- pixel layer, square- convolution layer, right-triangle- max-pooling layer, hexagon- fully connected layer, diamond- average pooling layer, down-triangle- local normalization layer, left-triangle- a skip module). Color changes from dark to light along the network.

##### 1.2.2 Manifold capacity's dependence on the number of objects and neurons

As discussed in the main text, while the theory is derived for a large number of objects $P$ and number of neurons $N$, meaningful results are achieved already at reasonable values.

Main figure 9c-d exhibits that measuring capacity using a finite number of objects shows only a small dependence on the number of objects used, and that already for $P \approx 50$ the measured capacity is a good approximation for the asymptotic value. Figure 13 shows similar results for 1-d and 2-d smooth manifolds, and using different levels of manifold variability.

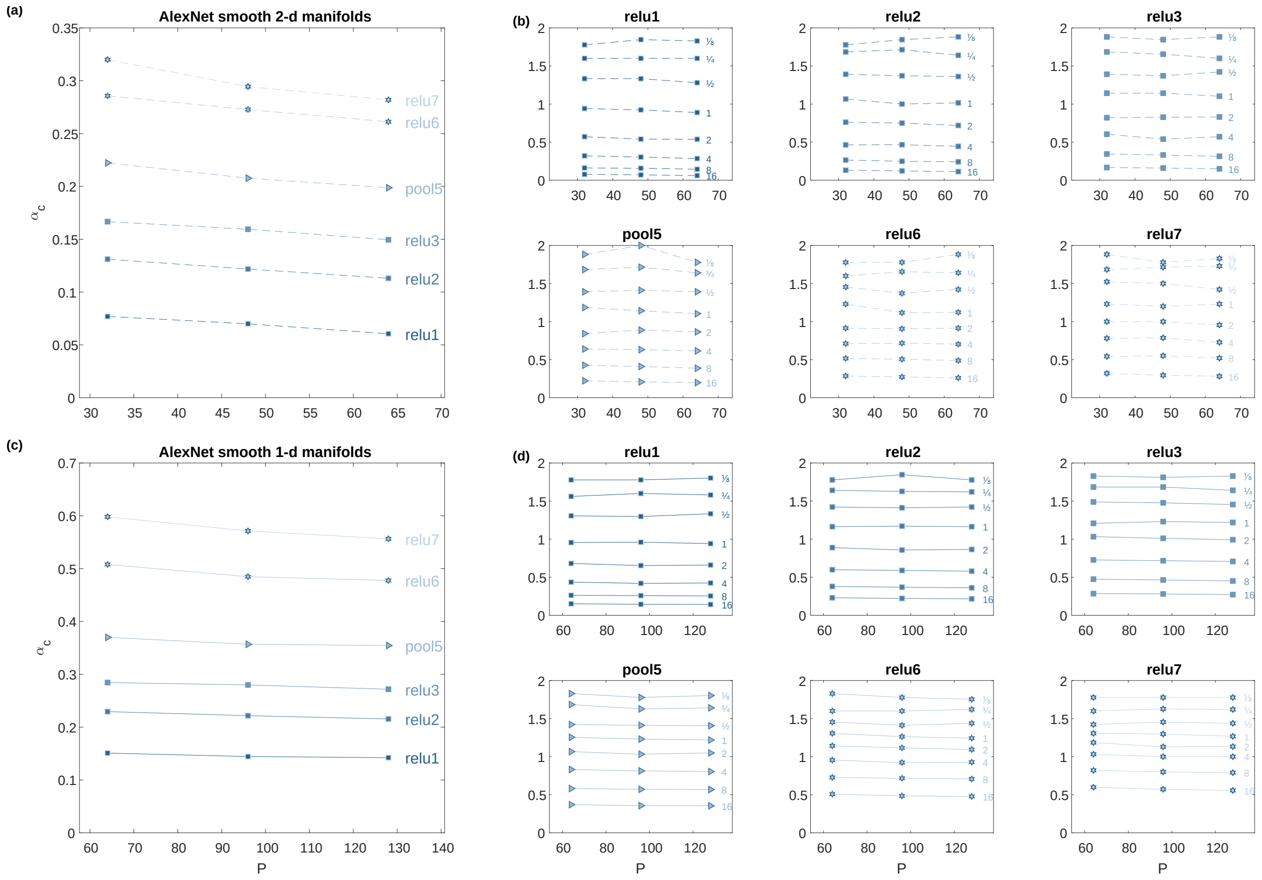

**Figure 13: Extensivity of numerically measured capacity.** Numerically measured capacity (y-axis) at different number of objects (x-axis) for smooth manifolds at 6 layers across AlexNet and 8 levels of variability. Results from 2-d shear manifolds at different layers at the largest variability level (a) and at 8 different levels of variability (indicated by the text, the maximal displacement of the object corners), one panel per layer (b). Results from 1-d translation manifolds at different layers at the largest variability level (c) and at 8 different levels of variability (indicated by the text, the maximal displacement of the object corners), one panel per layer (d). Marker shape represents layer type (circle- pixel layer, square- convolution layer, right-triangle- max-pooling layer, hexagon- fully connected layer). Color changes from dark to light along the network.

Similarly, when using mean-field algorithms to measure capacity and geometric properties using a finite number of neurons we expect the result to have only a small dependence on the number of neurons used. Figure 14 shows this is indeed the case; already when using a few hundred neurons, a good approximation of the asymptotic value is achieved.

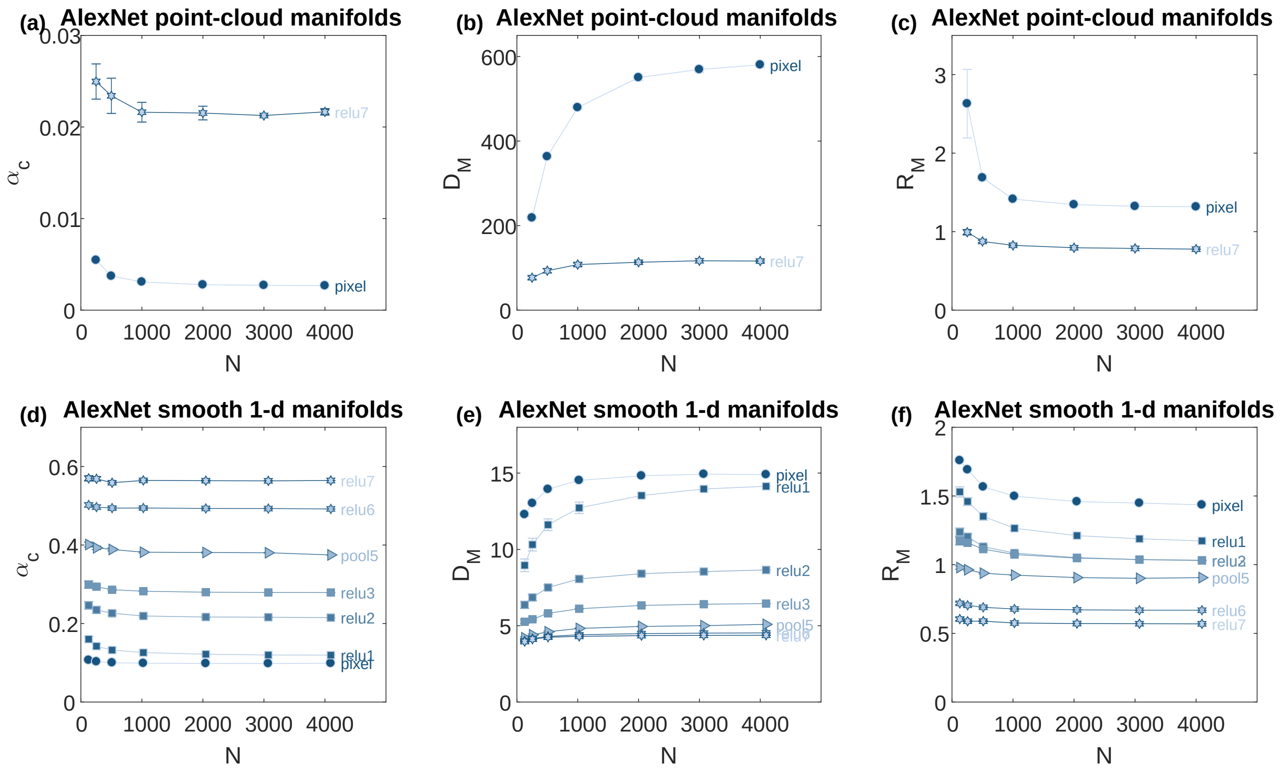

**Figure 14: Extensivity of capacity and manifold geometric properties.** Results from measuring manifolds classification capacity and geometric properties by subsampling different number of neurons; values are averaged across subsampling sets (5 for point-cloud manifolds, 10 for smooth manifolds); error-bars indicate standard deviation (and are of negligible size when not visible).
(a-c) Results for top 10% point-cloud manifolds (y-axis) vs. the number of subsampled neurons (x-axis): (a) classification capacity; (b) mean manifold dimension; (c) mean manifold radius.
(d-f) Results for smooth 1-d translation manifolds (y-axis) vs. the number of subsampled neurons (x-axis): (d) classification capacity; (e) mean manifold dimension; (f) mean manifold radius.
Marker shape represents layer type (circle- pixel layer, square- convolution layer, right-triangle- max-pooling layer, hexagon- fully connected layer). Color changes from dark to light along the network.

##### 1.2.3 Comparison between full theory and balls approximation for capacity in smooth manifolds of AlexNet, VGG-16, ResNet-50

Here we demonstrate that a manifold's classification capacity computed using the full mean-field theory (main text equation (1)) is well approximated by the classification capacity of $L_2$ balls with radius and dimension equivalent to the manifold's effective radius and dimension (computed from the distribution of anchor points, main text equations (3)-(5); see Methods). Thus $\alpha_c \approx \alpha_{Balls}(R_M, D_M)$ where the manifold capacity of $L_2$ balls is given by [2]:

$$\alpha_{\text{Ball}}^{-1}(R,D) = \int_{-\frac{\sqrt{D}}{R}}^{R\sqrt{D}} Dt_0 \frac{(R\sqrt{D}-t_0)^2}{R^2+1} + \int_{-\infty}^{-\frac{\sqrt{D}}{R}} Dt_0 (t_0^2 + D) \tag{4}$$

with Gaussian measure $Dt_0 = \frac{1}{\sqrt{2\pi}} e^{-\frac{t_0^2}{2}}$, which was first introduced in [3]. The agreement between $\alpha_c$ and $\alpha_{Balls}(R_M, D_M)$ is shown in various networks for smooth manifolds (figure 15).

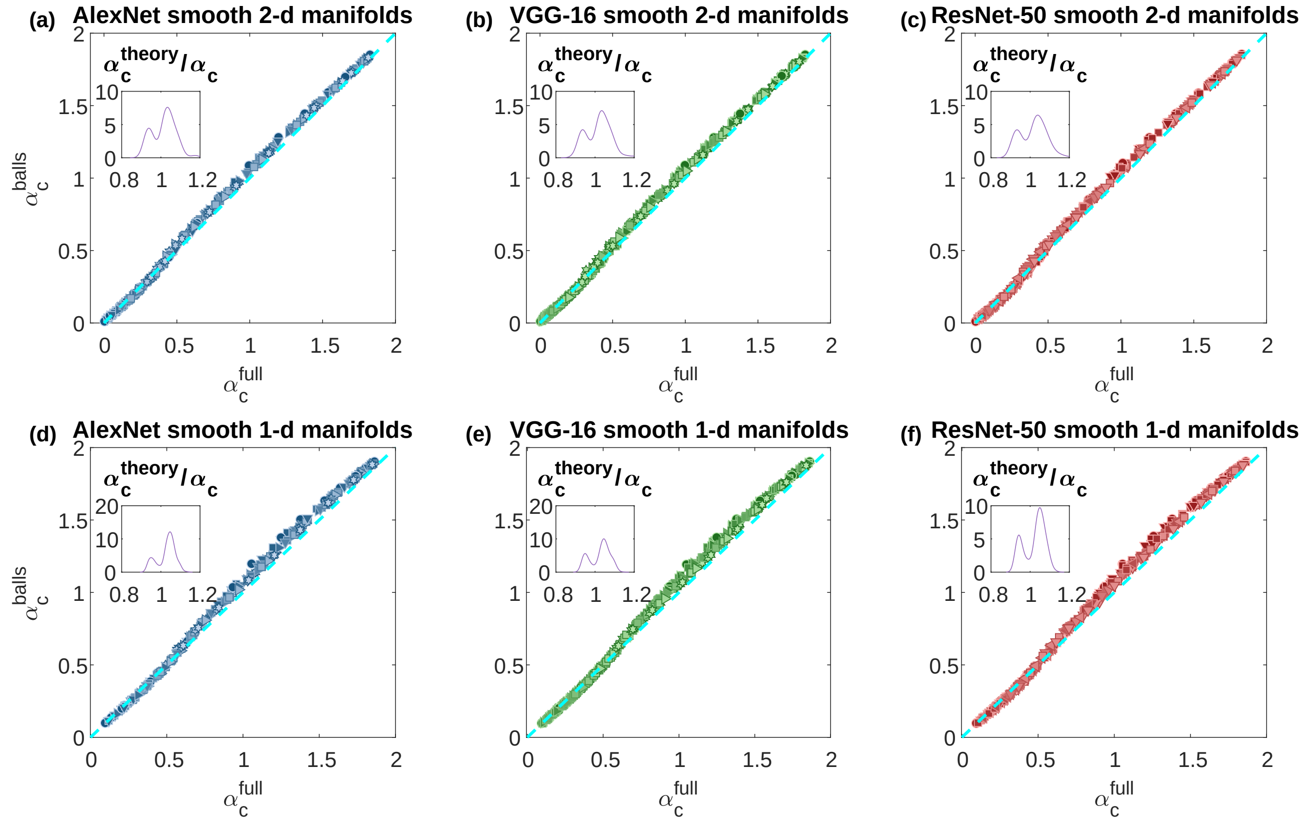

**Figure 15: Comparison of capacity measured using full theory and balls approximation.**
Comparison of capacity measured using the full theory (x-axis) with the approximation based on balls capacity and geometric properties (y-axis) for smooth manifolds at different layers along the hierarchy and different manifold variability levels. Inset: histogram of the ratio between the y-axis and the x-axis.
(a-c) Results for smooth 2-d manifolds across AlexNet (a), VGG-16 (b) and ResNet-50 (c).
(d-f) Results for smooth 1-d manifolds across AlexNet (d), VGG-16 (e) and ResNet-50 (f).
Marker shape represents layer type (circle- pixel layer, square- convolution layer, right-triangle- max-pooling layer, hexagon- fully connected layer, diamond- average pooling layer, down-triangle- local normalization layer, left-triangle- a skip module). Color changes from dark to light along the network.

##### 1.2.4 Random subsampling versus random projections

When numerically measuring capacity, the ability to linearly separate object manifolds is tested using a different number of neurons (see Methods in the main text). This requires to create representations of different dimensionality from an original high-dimensional representation; here, we compare two popular methods for doing so, random subsampling and using random projections. Comparisons of the numerically measured capacity using those sampling methods are shown in figure 16a and b for 1-d and 2-d smooth manifold, respectively. The good agreement between the two methods is demonstrated by the alignment of most points to the identity diagonal. A notable deviation from this agreement is evident in the above-diagonal square markers where capacity measured through subsampling is larger than that measured through random projections; those correspond to the 'relu5' layer where the neural response is very sparse. Figure 16c presents a sparsity histogram for the neurons of each layer, showing how 'relu5' is the only layer where the mode of this histogram is near 1.

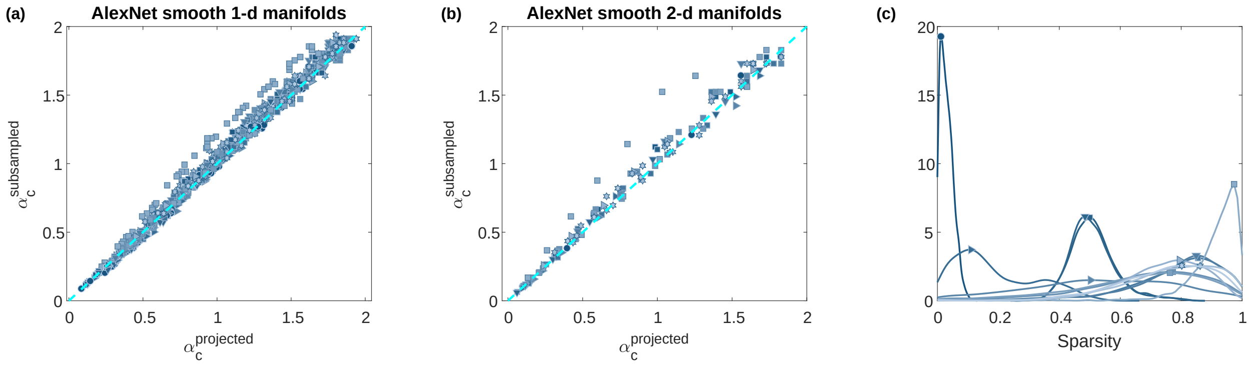

**Figure 16: Capacity using subsampled versus projected features.**
(a-b) Comparison of numerically measured capacity using random projections (x-axis) and using subsampling (y-axis) for AlexNet at different layers along the hierarchy and different levels of manifold variability.
(a) Smooth 1-d manifolds; (b) Smooth 2-d manifolds; (c) Histogram of per-neuron sparsity values for each layer; sparsity of a neuron is defined as the fraction of stimuli that elicit a zero response.
The identity diagonal is indicated by a dashed cyan line. Marker shape represents layer type (circle- pixel layer, square- convolution layer, right-triangle- max-pooling layer, hexagon- fully connected layer, down-triangle- local normalization layer). Color changes from dark to light along the network.

### 2 Methods

#### 2.1 Measuring capacity and geometric manifold properties

Here we develop a procedure to recover the common center correlations structure of general manifolds. Consider $P$ manifolds in $\mathbb{R}^N$ and denote $X \in \mathbb{R}^{N\times P}$ a matrix with their centers of mass; we seek to decompose the center correlations matrix $C = X^T X \in \mathbb{R}^{P\times P}$ into a sum of diagonal part $D$ and a low-rank part $C_K$ per main text equation (6). To achieve this, given $X$ we seek an orthonormal set $V \in \mathbb{R}^{N\times K}$ (i.e. $V^T V = I_K$) such that $X$ in the null-space of $V$ have approximately diagonal correlations:

$$\hat{X} = X - V\left(V^T X\right) \tag{5}$$

$$\hat{D} = \hat{X}^T \hat{X} \tag{6}$$

For specific $K$, this can be done by directly optimizing for $V$ which both satisfies the orthonormality constraint $V^T V = I_K$ and minimizes the following cost function (square_corrcoeff_cost() in Algorithm 1):

$$\text{cost}\left(X, V\right) = \frac{1}{2}\sum_{\mu\neq\nu} \hat{D}^2_{\mu\nu}/\hat{D}_{\mu\mu}\hat{D}_{\nu\nu} \tag{7}$$

where we note that $\hat{D}$ which minimizes this cost is expected to be approximately diagonal. This non-convex optimization problem can be efficiently solved using the algorithm and code from [4].

When $K$ is not known in advance, it can be found iteratively by taking increasing values for $K$, until the cost no longer decreases. To improve the stability of the algorithm we are using $V_K$ as the initial value for the optimization when optimizing for $V_{K+1}$; specifically we take an initial value of $[V_K|v]$ where $v$ is a randomly sampled vector and repeat the procedure several times to overcome dependencies on the initial conditions in this non-convex optimization. This approach achieves a minimal value for the cost at an intermediate value of $K$ so that it has no free parameter. Furthermore, the complexity of this procedure does not depend on the dimension $N$ as when $N \geq P$ the centers are of rank $P-1$ (as without loss of generality the global mean is 0) and thus can be expressed in $P-1$ coordinates. The full procedure for recovering the common components is described in Algorithm 1. Once the common components are removed, classification capacity, as well as manifold geometrical properties ($R_M$ and $D_M$, see main text), can be estimated from the residual manifolds using the methods described in [3]; the full procedure is described in Algorithm 2.

**Algorithm 1** find_common_components: find the number and directions of common components in correlated data

**Function** $V^*, X_K^*$ =find_common_components($X$)
**Input**: data $X \in \mathbb{R}^{N\times P}$
**Output**: common components matrix $V^* \in \mathbb{R}^{K\times N}$, data in the null-space of the common components $X_K^* \in \mathbb{R}^{N\times P}$
**Parameters**: maximal $K$ to try (a number smaller than $P$) and at the minima of which target function to stop (a boolean choosing mean square correlations or the mean absolute correlations)

1. Find an orthonormal set $Q \in \mathbb{R}^{N\times(P-1)}$ which spans the data, represent it in coordinates of $Q$ as $X_Q = Q^T X \in \mathbb{R}^{(P-1)\times P}$
2. Initialize $V = [\cdot]$ and the target function to $\infty$
3. Initialize $K = 0$, and until the maximal $K$ is reached or the target hasn't improved for 3 iterations:
   (a) Initialize $z = 0$
   (b) Repeat the optimization 10 times with different initial conditions to find the result with the smallest change to the norms of the data:
      i. Sample $\{s_\mu \sim \mathcal{N}(0,1)\}_{\mu=1}^{P}$ and set initial value $V_0 = [V|X_Q\vec{s}]$
      ii. Find $V_1$ which minimizes the square cost function under the constraint $V_1^T V_1 = I$ with initial value $V_0$: $V_1 = \text{OptStiefelGBB}(V_0, \text{square\_corrcoeff\_cost})$
      iii. Calculate residual data $\vec{x}_K^\mu = \vec{x}^\mu - V\left(V^T\vec{x}^\mu\right)$
      iv. Calculate $z_1 = \min_\mu \|\vec{x}_K^\mu\|/\|\vec{x}^\mu\|$
      v. If $z_1 > z$ save the results $V = V_1, z = z_1$
   (c) Calculate residual centers $X_K = X_Q - V\left(V^T X_Q\right)$
   (d) Calculate the target function (mean square correlations or mean absolute correlations)
   (e) If the current target function is lower than the best results so far, update $K^* = K$,$V^* = V$, $X_K^* = X_K$
4. Translate the results to original coordinates $V^* = QV^*$, $X_K^* = QX_K^*$

**Function** cost=square_corrcoeff_cost($V$,$X$)
**Input**: orthogonal matrix $V \in \mathbb{R}^{N\times K}$ and data matrix $X \in \mathbb{R}^{N\times P}$
**Output**: sum of squares of the off-diagonal normalized correlations between vectors of $X$ in the null-space of $V$

1. $C = X^T X$
2. $\hat{X} = X^T V$
3. $\hat{D}_{\mu\nu} = C_{\mu\nu} - \sum_i \hat{X}_{\mu i}\hat{X}_{\nu i}$
4. cost $= \frac{1}{2}\sum_{\mu\nu} \hat{D}_{\mu\nu}^2/\hat{D}_{\mu\mu}\hat{D}_{\nu\nu}$

**Function** $V$=OptStiefelGBB($V_0$, cost($\cdot$)): curvilinear search algorithm for optimization on Stiefel manifold
**Input**: an initial value $V_0 \in \mathbb{R}^{N\times K}$ and a cost function cost$(V)$
**Output**: $V \in \mathbb{R}^{N\times K}$ which minimizes cost$(V)$ under the constraint $V^T V = I$
See [4]

**Algorithm 2** correlated_manifolds_geometry: Geometric properties for correlated manifolds

**Function** correlated_manifolds_geometry($\{F^\mu\}$)
**Input**: object manifolds samples $\left\{F_i^\mu \in \mathbb{R}^N\right\}_{i\in[1..M_\mu]}^{\mu=1..P}$

1. Project data to the null-space of the common components: $\{f^\mu\} = \text{find\_residual\_data}\left(\{F^\mu\}\right)$
2. For $\mu = 1..P$, calculate geometry $D_M^\mu, R_M^\mu, \alpha_c^\mu = \text{manifold\_geometry}\left(f^\mu\right)$

**Output**: $\{D_M^\mu\}_{\mu=1}^P$, $\{R_M^\mu\}_{\mu=1}^P$, $\{\alpha_c^\mu\}_{\mu=1}^P$
**Function** $\{f^\mu\}$=find_residual_data($\{F^\mu\}$)
**Input**: object manifolds samples with $N$ neurons, $P$ objects, $M_\mu$ samples per object $\left\{F_i^\mu \in \mathbb{R}^N\right\}_{i=1..M_\mu}^{\mu=1..P}$
**Output**: residual object manifolds samples $\left\{f_i^\mu \in \mathbb{R}^N\right\}_{i=1..M_\mu}^{\mu=1..P}$

1. Remove the global mean from the input $F$
2. Calculate object centers $X \in \mathbb{R}^{N\times P}$
3. Find the common components $V = \text{find\_common\_components}\left(\text{X}\right)$
4. Use $V$ to find the object manifolds in the null-space of $V$: $f^\mu = F^\mu - V\left(V^T F^\mu\right)$

**Function** manifold_geometry($f^\mu$)
**Input**: object manifold $\left\{f_i^\mu \in \mathbb{R}^N\right\}_{i=1..M_\mu}$
**Parameters**: Numbers of Gaussian samples $N_G$
**Output**: manifold geometry $D_M^\mu$,$R_M^\mu$,$\alpha_c^\mu$
See [2] supplementary material, algorithms 2, 4.

#### 2.2 Measuring manifold capacity numerically

The details of how to numerically measure classification capacity, without making any of the assumptions of the mean-field theory, are provided in Algorithm 3.

**Algorithm 3** numerical_capacity: Measure numerical capacity

**Function**: numerical_capacity($\{F_i^\mu\}$)
**Input**: object manifolds samples with $N$ neurons, $P$ objects, $M_\mu$ samples per object $\left\{F_i^\mu \in \mathbb{R}^N\right\}_{i=1..M_\mu}^{\mu=1..P}$
**Output**: numerical capacity $\alpha_c$

1. Binary search for $n$ where fraction_separable_dichotomies $\left(\{F_i^\mu\}, n\right)$ surpass 0.5
2. Report $\alpha_c = P/n$

**Function** fraction_separable_dichotomies($\{F_i^\mu\}$, $n$): measure the fraction of separable random dichotomies
**Input**: object manifolds samples with $N$ neurons, $P$ objects, $M_\mu$ samples per object $\left\{F_i^\mu \in \mathbb{R}^N\right\}_{i=1..M_\mu}^{\mu=1..P}$ and an integer $n \in [0..N]$
**Output**: the fraction of linearly separable dichotomies after projecting the input to $n$
**Parameters**: $N_{dichotomies}$ the number of dichotomies to sample

1. Repeat $N_{dichotomies}$ times:
   (a) Sample random projections matrix $B \in \mathbb{R}^{n\times N}$ and set $f^\mu = BF^\mu\ \forall\mu = 1..P$
   (b) Sample random labeling $y \in \{\pm1\}^P$
   (c) Use quadratic optimization or the method from [5] to check if there exists $w \in \mathbb{R}^n$ such that $y^\mu w^T f_i^\mu \geq 0\ \forall\mu = 1..P\ \forall i = 1..M_\mu$
2. Report the fraction of separable dichotomies

#### 2.3 ImageNet classes used for point-cloud manifolds

The following ImageNet classes (a total of 50) were used as the first set of objects in the analysis of point-cloud manifolds.

moving van (n03796401)
hornbill (n01829413)
Tibetan terrier (n02097474)
microphone, mike (n03759954)
terrapin (n01667778)
wheelbarrow (n02797295)
race car (n04037443)
Weimaraner (n02092339)
knot (n03627232)
folding chair (n03376595)
sunscreen (n04357314)
coffeepot (n03063689)
overskirt (n03866082)
fox squirrel (n02356798)
standard poodle (n02113799)
oscilloscope (n03857828)
head cabbage (n07714571)
jeweler's loupe (n03692522)
bottlecap (n02877765)
llama (n02437616)
ruddy turnstone (n02025239)
hermit crab (n01986214)
ibex (n02417914)
vase (n04522168)
little blue heron (n02009229)
soda bottle (n03983396)
rotisserie (n04111531)
microwave oven (n03761084)
breastplate (n02895154)
home theater (n03529860)
black widow (n01774384)
mailbox (n03710193)
crutch (n03141823)
bighorn sheep (n02415577)
hartebeest (n02422106)
padlock (n03874599)
sewing machine (n04179913)
mountain bike (n03792782)
giant panda (n02510455)
flowerpot (n03991062)
bow (n02879718)
triceratops (n01704323)
otter hound (n02091635)
chain (n02999410)
cowboy hat (n03124170)
tripod (n04485082)
bell (n03017168)
smoothing iron (n03584829)
teapot (n04398044)
birdhouse (n02843684)

### 3 Notes

#### 3.1 Theory for low-rank center correlations

**Uncorrelated general manifolds** Consider $P$ manifolds $\{M^\mu\}_{\mu=1}^P$ which reside in $\mathbb{R}^N$ but have an intrinsic dimension of $D+1$:

$$M^\mu = \left\{ \sum_{l=0}^{D} s_l \vec{u}^{\mu,l} \ : \ \vec{s} \in S^\mu \right\} \tag{8}$$

where $S^\mu \subseteq \mathbb{R}^{D+1}$ are the manifold coordinates and the unit vectors $\left\{\vec{u}^{\mu,l} \in \mathbb{R}^N\right\}_{\mu=1..P}^{l=0..D}$ are the axes on which the (non-linear) manifold is defined. By convention $\vec{u}^{\mu,0}$ is the manifold center and $s_0$ is its norm in this parametrization. The manifold coordinates are defined by a set of functions $F_\mu(x) : \mathbb{R}^{D+1} \to \mathbb{R}$ for $\mu = 1..P$:

$$S^\mu = \{s \ : \ F_\mu(s) < 0\} \tag{9}$$

Denote $\{y^\mu\}_{\mu=1}^P$ random labels and $\alpha = \frac{P}{N}$ the system's load, we are interested in the case $N, P \to \infty$ with a finite $\alpha$ and ask when there exists a solution, that is a unit vector $\vec{w} \in \mathbb{R}^N$, a bias term $b \in \mathbb{R}$ such that:

$$y^\mu \left(w^T x + b\right) \geq 0 \ \forall x \in M^\mu \ \forall \mu = 1..P \tag{10}$$

The critical capacity $\alpha_c$ is the load value such that below it a solution is likely to exist and above it a solution is unlikely. Formally we consider the volume of the solution space under constraints of normalization of $w$ (which we take as $\|w\|^2 = 1$) and satisfaction of the solution inequalities:

$$V[\alpha] = \int db \prod_i^N \int dw_i \delta\left(w^T w - 1\right) \prod_\mu^P \prod_{x \in M^\mu} \Theta\left(y^\mu \left(w^T x + b\right)\right) \tag{11}$$

where $\Theta$ is the Heaviside step function.

By assuming $\{u_i^{\mu,l}\}_{i=1..N}^{\mu=1..P,l=0..D}$ are i.i.d standard Gaussian variables $\mathcal{N}(0,1)$, thus avoiding any correlations between manifolds axes of variation and between manifold centers, and considering binary labeling $y^\mu \in \{\pm 1\}$ with a equal probability, it can be shown that the critical capacity satisfies [2], for homogeneous manifolds:

$$\alpha_S^{-1} \ = \ \int D^{D+1}\vec{t} \min_{\{\vec{v} \in \mathbb{R}^{D+1} \, : \, \forall \vec{s} \in S \ \vec{s} \cdot \vec{v}^\mu \geq 0\}} \|\vec{v}^\mu - \vec{t}^\mu\|^2 \tag{12}$$

where $Dt = dt e^{-t^2/2}/\sqrt{2\pi}$ and for non-homogeneous manifolds:

$$\alpha_c^{-1} \ = \ \frac{1}{P} \sum_\mu^P \alpha_{S^\mu}^{-1} \tag{13}$$

**Manifolds with full-rank center correlations** Consider the more general case where the manifold center (or displacement relative to the origin) is correlated across manifolds, that is $\{\vec{u}^{\mu,0} \in \mathbb{R}^N\}_{\mu=1}^P$ have correlations. Denote those correlations:

$$C_{\mu\nu} \ = \ \frac{1}{N} \left[\vec{u}^{\mu,0} \cdot \vec{u}^{\nu,0}\right]_u \tag{14}$$

Assuming the correlations are of full rank, there exist unit vectors $\{\vec{r}^l \in \mathbb{R}^P\}_{l=1}^P$, and scalars $\{c_l \geq 0\}_{l=1}^P$ such that:

$$C_{\mu\nu} \ = \ \sum_l^P c_l r_\mu^l r_\nu^l \tag{15}$$

and denoting a matrix $L_{\mu\nu} = \sqrt{c_\nu} \vec{r}_\mu^\nu$ we have that:

$$C \ = \ \sum_l^P c_l \vec{r}^l \vec{r}^{lT} = LL^T \tag{16}$$

Now using $\{\phi_{l,i} \sim \mathcal{N}(0,1)\}_{l=1..P}^{i=1..N}$ i.i.d Gaussian variables we define a statistical model:

$$u_i^{\mu,0} = \sum_l^P \sqrt{c_l} r_\mu^l \phi_{l,i} \tag{17}$$

which satisfies supplementary equation (14).

Following [6],[2] the critical capacity is characterized by the volume of solutions using the replica method; assuming replica symmetry we get an expression:

$$\alpha_c^{-1} = \int D^{(D+1)\times P} t_{l=0..D}^{\mu=1..P} \left[ \min_{\vec{v}\in\mathcal{V}} \frac{1}{P} \sum_\mu^P \|\vec{v}^\mu - \vec{t}^\mu\|^2 \right]_y \tag{18}$$

$$\mathcal{V} = \left\{ \vec{v} \in \mathbb{R}^{(D+1)\times P} \; : \; \forall \mu \, \forall \vec{s}^\mu \in S^\mu \; \sum_{l=1}^D s_l v_l^\mu + y^\mu \sum_\nu L_{\mu\nu} v_0^\nu \geq 0 \right\} \tag{19}$$

where the matrix $L$ couples the integration between different manifolds and the expression still depends on the labeling $\vec{y}$; compare to supplementary equation (13) from of the uncorrelated case written using a similar notation:

$$\alpha_c^{-1} = \int D^{(D+1)\times P} t_l^\mu \min_{\vec{v}\in\mathcal{V}} \frac{1}{P} \sum_\mu^P \|\vec{v}^\mu - \vec{t}^\mu\|^2 \tag{20}$$

$$\mathcal{V} = \left\{ \vec{v} \in \mathbb{R}^{(D+1)\times P} \; : \; \forall \mu \; \forall \vec{s}^\mu \in S^\mu \; \vec{s}^\mu \cdot \vec{v}^\mu \geq 0 \right\} \tag{21}$$

**Manifolds with low-rank off-diagonal correlations** For correlations with low-rank off-diagonal structure we assume there is a diagonal $\Delta$ with $\Delta_{\mu\mu} = d_\mu$ and $C_K$ with rank $K \ll P$ such that:

$$C = \Delta + C_K \tag{22}$$

Noting that $\Delta^{-0.5} C_K \Delta^{-0.5}$ is symmetric there exists an orthonormal $U_K \in \mathbb{R}^{P\times K}$ and a diagonal $E_K \in \mathbb{R}^{K\times K}$ such that:

$$U_K E_K U_K^T = \Delta^{-0.5} C_K \Delta^{-0.5} \tag{23}$$

Denote $U$ a completion of $U_K$ to orthonormal basis and denote $E \in \mathbb{R}^{P\times P}$ defined as $E = \begin{bmatrix} E_K & 0 \\ 0 & 0 \end{bmatrix}$, we have:

$$C = \Delta^{0.5} \left( I + U_K E_K U_K^T \right) \Delta^{0.5} = \Delta^{0.5} U \left( I + E \right) U^T \Delta^{0.5} \tag{24}$$

so that $C = LL^T$ for:

$$L = \Delta^{0.5} U \left( I + E \right)^{0.5} \tag{25}$$

Using this notation capacity from supplementary equation (20) becomes:

$$\alpha_c^{-1} = \int D^{(D+1)\times P} t_l^\mu \left[ \min_{\vec{v}\in\mathcal{V}} \left[ \frac{1}{P} \sum_{l=0}^D \|\vec{v}_l - \vec{t}_l\|^2 \right] \right]_y \tag{26}$$

$$\mathcal{V} = \left\{ \vec{v} \in \mathbb{R}^{(D+1)\times P} \; : \; \forall \mu \, \forall \vec{s}^\mu \in S^\mu \; \sum_{l=1}^D s_l^\mu v_l^\mu + y^\mu \sqrt{d_\mu} \sum_\nu U_{\mu\nu} \sqrt{1+e_\nu} v_0^\nu \geq 0 \right\} \tag{27}$$

Now denote $\alpha_{ns}$ an approximation assuming the minima with respect to $\vec{v}_0$ is achieved for $v_0^\nu = 0$ for $\nu = 1..K$, and further neglect the contribution of the corresponding $t_0^\nu$; as long as $K \ll P$ we can expect this approximation to be reasonable, yielding:

$$\alpha_{ns}^{-1} = \int D^{(D+1)\times P} t_l^\mu \left[ \min_{\vec{v}\in\mathcal{V}} \left[ \frac{1}{P} \sum_{l=1}^D \|\vec{v}_l - \vec{t}_l\|^2 + \frac{1}{P} \|\vec{v}_0 - t_0\|^2 \right] \right]_y \tag{28}$$

$$\mathcal{V} = \left\{ \vec{v} \in \mathbb{R}^{(D+1)\times P} \; : \; \forall \mu \, \forall \vec{s}^\mu \in S^\mu \; \sum_{l=1}^D s_l^\mu v_l^\mu + \sqrt{d_\mu} y^\mu \sum_{\nu=K+1}^P U_{\mu\nu} v_0^\nu \geq 0 \right\} \tag{29}$$

such that by change of variables $\vec{v}_0 \leftarrow \vec{y} \circ U \vec{v}_0$ and $\vec{t}_0 \leftarrow \vec{y} \circ U \vec{t}_0$ which does not affect the norm $\|\vec{v}_0 - t_0\|^2$ we have a decoupled expression:

$$\alpha_{ns}^{-1} = \int D^{(D+1)\times P} t_l^\mu \min_{\vec{v}\in\mathcal{V}} \left[\frac{1}{P}\sum_{l=0}^{D} \|\vec{v}_l - \vec{t}_l\|^2\right] \tag{30}$$

$$\mathcal{V} = \left\{\vec{v}\in\mathbb{R}^{(D+1)\times P} \; : \; \forall\mu\, \forall\vec{s}^\mu \in S^\mu \; \min_{\vec{s}\in S^\mu} \sum_{l=1}^{D} s_l v_l^\mu + \sqrt{d_\mu} v_0^\mu \geq 0\right\} \tag{31}$$

Thus by projecting the manifolds into the null-space of the directions associated with the non-diagonal part of the correlations matrix, we are back at the situation of manifolds with uncorrelated centers and thus can use capacity from supplementary equation (13) with appropriate scaling of the manifolds by the norm $\{d_\mu\}$.

##### 3.2 Theory for manifolds of random points

**Properties of point-cloud manifolds** Consider a point-cloud manifold composed of $M$ points $F = \left\{\vec{x}^m \in \mathbb{R}^N\right\}_{m=1}^M$ where $N$ denotes the ambient dimension and assume the points reside in a $D+1$ affine subspace, such that there is an orthonormal set $\left\{u_l \in \mathbb{R}^N\right\}_{l=0}^D$ such that:

$$\vec{x}^m = s_0\vec{u}_0 + \sum_{l=1}^{D} s_l^m \vec{u}_l \tag{32}$$

where $\left\{\vec{s}^m \in \mathbb{R}^D\right\}_{m=1}^M$ denote manifold coordinates and $s_0$ denotes the manifold's center norm.
In this setup classification capacity is given by:

$$\alpha_c^{-1} = \int D^{D+1}\vec{t} \min_{S\vec{v}\geq 0} \|\vec{v} - \vec{t}\|^2 \tag{33}$$

where $S \in \mathbb{R}^{M\times(D+1)}$ denotes the manifold coordinates such that the $D+1$ coordinate of each row represents the manifold center:

$$S = \begin{bmatrix} \vec{s}^1 - \bar{s} & s_0 \\ \vdots & s_0 \\ \vec{s}^m - \bar{s} & s_0 \end{bmatrix} \tag{34}$$

$$\bar{s} = \frac{1}{M}\sum \vec{s}^m \tag{35}$$

$$s_0 = \|\bar{s}\| \tag{36}$$

Thus exact calculation of capacity requires to solve the following optimization problem:

$$\vec{v}\left(\vec{t}\right) = \arg\min_{S\vec{v}\geq 0} \|\vec{v} - \vec{t}\|^2 \tag{37}$$

with Lagrangian (written such that the $D+1$ coordinate is denoted $v_0$, $t_0$):

$$\mathcal{L} = \frac{1}{2}\sum_{l=0}^{D}(v_l - t_l)^2 - \sum_{m}^{M}\lambda_m\left(\sum_{l=0}^{D} s_l^m v_l - \sum_{l=0}^{D}\bar{s}_l v_l + s_0 v_0\right) \tag{38}$$

Then manifold properties are defined in terms of the solution:

$$\tilde{s}_l = \frac{v_l - t_l}{v_0 - t_0} \tag{39}$$

$$R_M = \left[\sqrt{\sum_{l}^{D}\delta\tilde{s}_l^2}\right] \tag{40}$$

$$D_M = \left[\sum_{l}^{D}\frac{t_l\delta\tilde{s}_l}{\|\delta\tilde{s}\|}\right]^2 \tag{41}$$

**Properties of random point-cloud manifolds** We seek to analyze $R_M$ and $D_M$ for the case of random point-cloud manifolds, i.e. when $\{s_l^m \sim \mathcal{N}(0,1)\}_{m=1..M}^{l=0..D}$ and are i.i.d sampled. When $D \gg M$ for a random $\{\vec{s}^m\}_{m=1}^M$ we further assume that $\vec{s}^{m_1} \perp \vec{s}^{m_2}$ for $m_1 \neq m_2$ and thus can simplify the analysis through an orthogonal change of basis from $D$ to the standard basis $\vec{s}^m = \vec{e}^m$ of an $M$-dimensional space. In this case $\bar{s} \equiv 1/M$ so that $s_0 = 1/\sqrt{M}$:

$$S = \begin{bmatrix} \vec{e}^1 - 1/M & s_0 \\ \vdots & s_0 \\ \vec{e}^m - 1/M & s_0 \end{bmatrix} \tag{42}$$

so the Lagrangian becomes:

$$\mathcal{L} = \frac{1}{2}\sum_i^M (v_i - t_i)^2 + \frac{1}{2}(v_0 - t_0)^2 - \sum_m^M \lambda_m \left( v_m - \frac{1}{M}\sum_j^M v_j + s_0 v_0 \right) \tag{43}$$

An exact solution to this optimization problem is derived using Kuhn-Tucker conditions and is given by:

$$\lambda_l\left(\vec{t}\right) = \left[ -t_l + \frac{1}{M}\sum_j^M t_j - \frac{t_0}{\sqrt{M}} \right]_+ \tag{44}$$

$$\bar{\lambda}\left(\vec{t}\right) = \frac{1}{M}\sum \lambda_l \tag{45}$$

$$v_l\left(\vec{t}\right) = t_l + \lambda_l - \bar{\lambda} \tag{46}$$

$$v_0\left(\vec{t}\right) = t_0 + \sqrt{M}\bar{\lambda} \tag{47}$$

Then the exact expression for $\delta\vec{s}$ is:

$$\delta\tilde{s}_l = \frac{v_l - t_l}{v_0 - t_0} = \frac{\lambda_l - \bar{\lambda}}{\bar{\lambda}\sqrt{M}} \tag{48}$$

By replacing the exact expression for $\lambda$ from equation (44) with an approximation:

$$\lambda_l\left(\vec{t}\right) = \left[-t_l\right]_+ \tag{49}$$

we can analytically calculate the mean-field manifold properties:

$$D_M = \frac{\pi}{2(\pi - 1)} M \tag{50}$$

$$R_M = \sqrt{\pi - 1} \tag{51}$$

$$\alpha_c = \frac{2}{M} \tag{52}$$
